# Combined stimuli of elasticity and microgrooves form aligned myotubes that characterize slow-twitch muscles

**DOI:** 10.1101/2025.03.30.646214

**Authors:** Hiroki Hamaguchi, Tomoko G. Oyama, Kotaro Oyama, Yasuko Manabe, Nobuharu L. Fujii, Mitsumasa Taguchi

## Abstract

Skeletal muscles are classified into slow-twitch muscles composed primarily of type I and IIa fibers with high oxidative metabolism, and fast-twitch muscles composed of type IIx and IIb fibers with high glycolytic metabolism. Fiber-type shifts occur during development and aging; however, the stimuli that shift these types remain unclear. We analyzed the role of mechanical stimuli in myotube formation and shift to the characteristics of each fiber type using crosslinked gelatin gels with tunable elasticity (10–230 kPa) and microgrooves (3–50 µm). C2C12 myotubes on 10 kPa gel increased the expression of marker genes for type I and IIa fibers *(MYH7* and *MYH2*) and oxidative metabolism (*GLUT4* and *myoglobin*) than those on stiffer gels. Upregulation of *PGC-1α* on soft gel induced a shift toward slow-twitch muscle genetic characteristics. Microgrooves (3–10 µm) enhanced myoblast fusion and myotube orientation, without affecting the gene expressions characterizing fiber types. This study demonstrated an approach to create highly oriented slow-twitch muscle models by controlling the elasticity and microgrooves.

## Introduction

Skeletal muscle is the largest tissue in the human body and is essential for physical exercise and metabolism [1, 2]. It is broadly classified into two types, slow and fast-twitch muscles, based on differences in contraction and metabolic characteristics [3]. Slow-twitch muscles, such as the soleus, cannot exert great force but exert constant force for long periods during activities, such as walking, postural control, and endurance exercise. Additionally, these muscles have high glucose tolerance, enabling them to absorb glucose from the blood and maintain normal blood glucose levels. Moreover, they exhibit a high oxidative metabolism capacity to synthesize ATP through the mitochondrial electron transfer system. In contrast, fast-twitch muscles, such as the extensor digitorum longus, have low endurance but are active in movements and exercises that require instantaneous force. Additionally, these muscles are characterized by high glycolytic metabolism, where synthesizing ATP through the degradation of glucose to pyruvate. These differences in characteristics depend on the type of muscle fibers (i.e., mature skeletal muscle cells) that constitute the skeletal muscle. There are four types of myofibers: I, IIa, IIx, and IIb, and each type is determined by the expression level of the *myosin heavy chain* (*MYH*) isoforms—*MYH7*, *MYH2*, *MYH1*, and *MYH4*—that regulate muscle contraction. Type I and IIa fibers have high oxidative metabolism, whereas type IIx and IIb fibers have high glycolytic metabolism [3]. Therefore, skeletal muscles composed primarily of type I and IIa fibers exhibit slow-twitch muscle characteristics, whereas those composed of type IIx and IIb fibers exhibit fast-twitch muscle characteristics.

During development, myofiber types of skeletal muscle cells undergo considerable changes [3, 4]. In the early stages of myogenesis, *MYH7*, a marker of type I fibers, is initially expressed. Subsequently, *MYH* isoform expression diversifies, leading to fiber type diversification. As skeletal muscle matures, neural activity and mechanical loading facilitate a shift from type I to type IIa, IIx, and IIb fibers. The type and composition of muscle fibers are determined by species, body parts, and genetics [5,6,7,8]. Over time, chronic exercise training alters the distribution of *MYH* isoforms in muscles, increasing the proportion of *MYH7* and/or *MYH2* [9]. Moreover, it enhances mitochondrial neogenesis and glucose metabolism in myofibers and induces a shift from type IIx and IIb fibers to type I and IIa [10]. However, the specific stimuli that induce a shift to each fiber type during development and later in life remain unclear. Additionally, myofiber types differ in their susceptibility to muscle atrophy induced by aging, inactivity, diabetes, and cancer [11, 12]. Therefore, if the stimuli and mechanisms that induce a shift to each fiber type are identified, strategies to increase specific fiber types can be applied to enhance exercise function and metabolic capacity, thereby enhancing health. Specifically, skeletal muscle is a major determinant of the basal metabolic rate and responsible for over 80 % of insulin-stimulated whole-body glucose uptake [13, 14]. If type I and IIa fibers with high oxidative metabolism increase, they may enhance whole-body metabolism by improving metabolic abnormalities such as insulin resistance.

Traditionally, experimental animals have been widely used to study fiber-type development and acquired shifts. However, analyzing myogenesis and fiber-type shifts in mature muscle tissues requires dissecting mice immediately after birth or when they are mature. Because this tissue is lined with nerves and capillaries, confounding factors, such as nerve-derived stimuli and humoral factors are included. Therefore, analyzing the stimuli and mechanisms that directly control fiber types is challenging, along with the cost, time, and ethics of the experiments. In skeletal muscle studies, myotube-differentiated skeletal muscle culture cells are frequently used instead of animal tissues. In vitro, myoblasts are seeded onto plastic (PS) dishes coated with extracellular matrix (ECM) extracts, such as collagen and laminin, and are subsequently differentiated into myotubes. However, myotubes exhibit disordered *MYH* isoform gene expression associated with the contractile properties of myofibers and metabolic-related genes and failed to reproduce the developmental process of skeletal muscle, as characterized by fiber type [15]. Sustained muscle contractions through electrical stimulation to mimic chronic training enhance the expression of *MYH7* and *MYH2* that are associated with contractile properties with higher expression levels in myofibers with predominant oxidative metabolism [16]. However, no alterations were induced in genes associated with oxidative metabolism. It is challenging to establish stimuli that induce both contractile and metabolic activities characteristic of each fiber type because of the possibility of cell damage or detachment during electrical stimulation. Therefore, it is significant to establish novel stimuli to form muscle models composed of each fiber type with contractile and metabolic properties without cell damage or detachment.

Cellular activities *in vivo* are influenced by the physical properties of the surrounding microenvironment (i.e., cell niche). Mechanical stimuli, such as elasticity and microgrooves from culture substrates, affect various cellular responses, including cell morphogenesis, migration, proliferation, apoptosis, and differentiation [17–21]. In skeletal muscle cells, the elasticity and microgrooves of the culture substrate induce myotube differentiation, maturation, and high orientation [22–30]. For example, Engler et al. demonstrated that the optimal elastic modulus for myotube differentiation is 12 kPa using polyacrylamide gels with striped patterns of crosslinked collagen [27]. Additionally, microgrooved gels enhance the orientation and myogenic differentiation of myotubes [22]. These studies indicate that mechanical stimuli derived from the elasticity and microgrooves of the culture substrate contribute to the development of skeletal muscle cells. However, to our knowledge, it has not been confirmed whether these mechanical stimuli contribute to the induction of shifts to each fiber type.

We hypothesized that the elasticity and microtopography of the culture substrate can contribute to the induction of myotube shifts to each fiber type. Skeletal muscle elasticity ranges from approximately 10–100 kPa, based on the region and literature [31–35]. Therefore, to assess this hypothesis, we required a culture substrate with tunable elasticity comparable to the physiological range of skeletal muscle and the potential to incorporate microtopography. Previously, we developed a technique for crosslinking and gelation of various proteins using radiation-induced reactions [36]. This technology controls the elasticity of protein gels by adjusting the protein concentration in an aqueous solution and irradiation dose. Combined with imprinting technology, microtopography is imparted to the surface during gelation. In this study, we applied this technology to gelatin, a highly purified form of type I collagen abundant in the ECM of skeletal muscle. Although myotubes do not differentiate into myofibers *in vitro*, the induction of fiber-type shifts in skeletal muscle cells can be assessed by evaluating the expression levels of the *MYH* isoforms and metabolic-related genes in myotubes. Therefore, we comprehensively analyzed whether mechanical stimuli from the elasticity and microgrooves of gelatin gels induced the genetic characteristics of each fiber type in C2C12 myotubes. The results revealed that the elasticity of gelatin gels induced a shift of myotubes to the genetic characteristics of slow-twitch muscles composed of type I and IIa fibers with high oxidative metabolism, whereas microgrooves enhanced myoblast fusion and myotube orientation without inducing the shift to the characteristics of any fiber types.

## Results

### Preparation of a gelatin gel with controlled elasticity and surface microgrooves

Crosslinked gelatin gels with controlled elasticity and surface microgrooves were prepared to assess the effect of mechanical stimulation on shifts to distinct fiber types in vitro (Fig. 1a). Type I collagen is the predominant collagen in the ECM of muscle tissue in vivo. However, we selected non-fibrous gelatin—a hydrolysate of type I collagen—to eliminate the effect of randomly formed collagen fibers and assess the effect of mechanical stimulation from artificially added microgroove shapes. The gelatin solution was irradiated with γ-rays to generate OH radicals from water radiolysis, to introduce crosslinks between the gelatin molecules. Gelatin is transformed into a three-dimensional mesh structure that encapsulates the surrounding water—crosslinked hydrogel. Unlike physical gels, the crosslinked gel did not dissolve even at 37 °C. This study prepared gels with compressive elastic moduli of approximately 10, 30, 50, 100, 160, and 230 kPa using a combination of aqueous solution concentrations (10 and 15 wt%) and γ-ray irradiation doses (8–30 kGy [= J/g]) to cover the reported elastic moduli of skeletal muscle (10–100 kPa) (Fig. 1b). By imprinting microstructures from a polydimethylsiloxane (PDMS) mold during gelation, microgrooves with width and spacing of 3, 5, 10, and 50 μm and a depth of 2 μm were fabricated on gels with varying elasticity. As a representative example, microgrooves formed on the surface of a 30 kPa gel are illustrated in Fig. 1c.

**Fig. 1.**
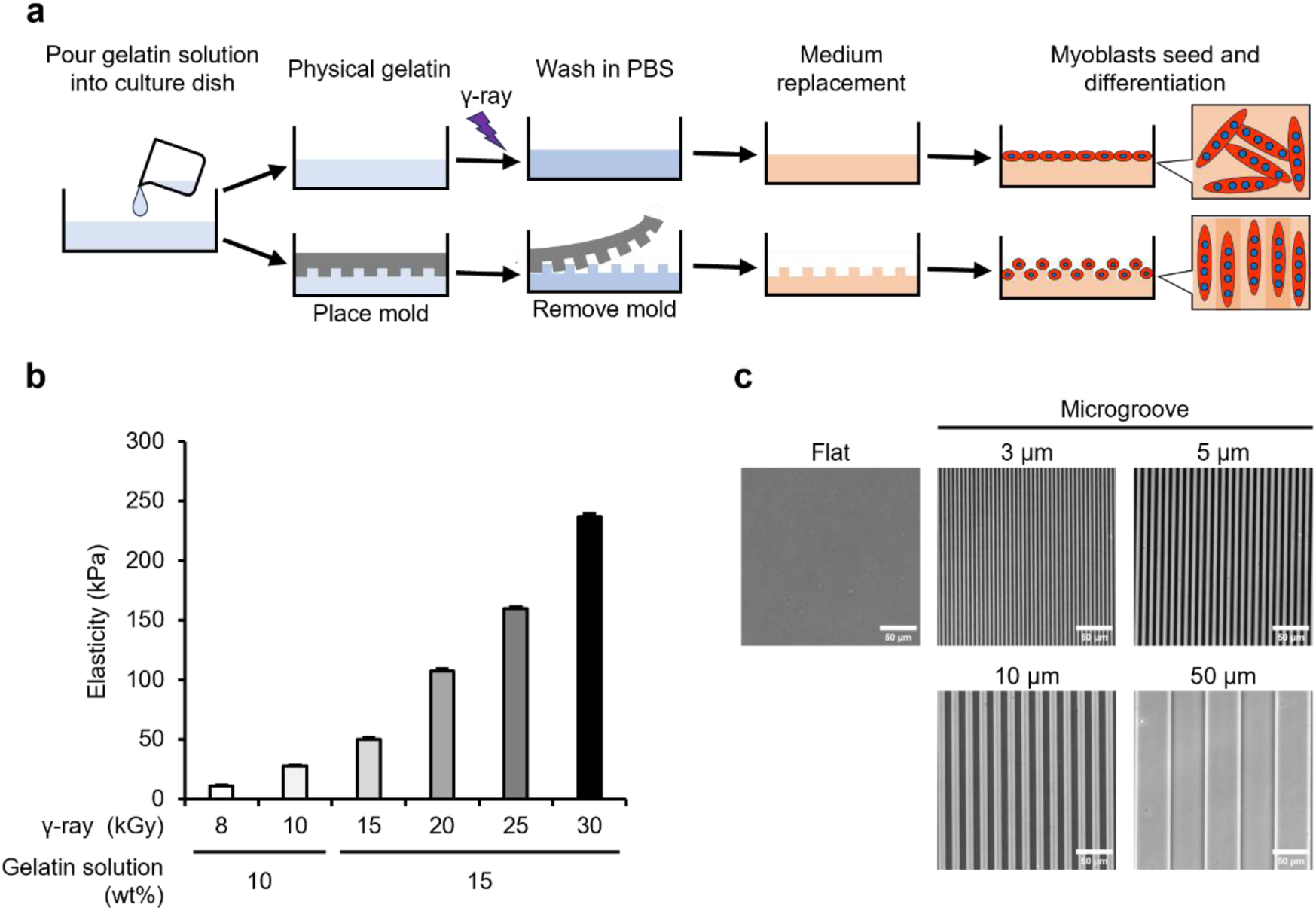
Preparation process, elasticity, and microgrooves of gelatin gels. (a) Preparation process of crosslinked gelatin gels using radiation-induced reaction. Gelatin physical gel is irradiated with γ-rays to induce crosslinking. The surface microgrooves are transferred from polydimethylsiloxane molds, and the gels are used for cell culture experiments after the phosphate-buffered saline (PBS) wash and medium replacement. (b) Compressive elastic modulus of gelatin gels prepared from combinations of concentration of gelatin solution (wt%) and γ-ray irradiation dose (kGy). Elasticity is measured using a rheometer. Data are presented as the mean ± standard error of the mean (S.E.M.), n = 3. (c) Phase contrast images of microgroove formed on the gelatin gel. The images demonstrate microgrooves on the 30 kPa gels prepared using an irradiation dose of 10 kGy. Scale bars, 50 μm.

### Adhesion, viability, proliferative rate, and area of myoblasts on gelatin gels

C2C12 myoblasts were seeded on flat gels (elasticity: 10–230 kPa) and PS dishes (commonly used for myoblast culture). The elasticity of PS dish is > 1 GPa, which is much stiffer than that of the gelatin gel that mimics the elasticity in vivo. Myoblasts adhered to gels of each elasticity and PS dish, and over 99 % of myoblasts survived under all conditions (Fig. 2a). Proliferative rate exhibited no difference between the gels of each elasticity and PS dishes at 24–48 h after seeding (Sup. Fig. 1a). In contrast, the myoblast area was significantly reduced on gels of all moduli compared to that on the PS dish and was smaller on gels of lower moduli (Sup. Fig. 1b). This difference in area affects the confluency (area fraction occupied by cells)—a crucial parameter for subsequent differentiation into myotubes. Therefore, in subsequent experiments, myoblasts were seeded at a uniform density of 4.2×10^4^ cells/cm^2^ to achieve approximately 80 % confluency, and differentiation was induced the following day.

**Fig. 2.**
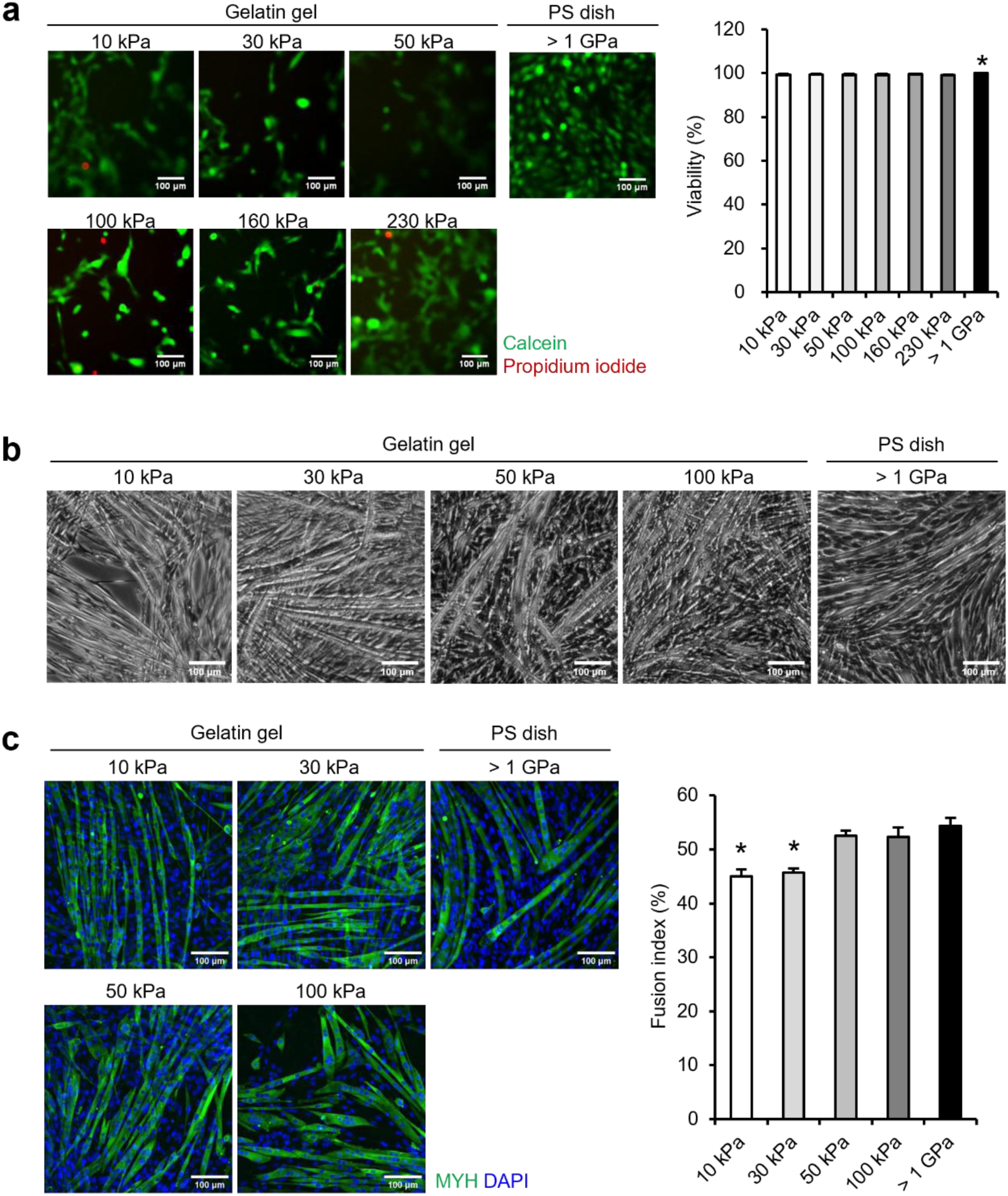
Viability of myoblasts on gelatin gel and differentiation into myotubes. (a) Viability of myoblasts cultured on the 10–230 kPa gelatin gels and plastic dishes (PS dish, > 1 GPa). Images demonstrate myoblasts at 24 h post-seeding stained with live cells (green: calcein) and dead cells (red: propidium iodide). Viability is calculated by dividing the number of live cells by the total number of cells (live cells + dead cells). (b) Phase contrast images of myotubes differentiated and cultured on 10–100 kPa gelatin gels and PS dishes. The images demonstrate myotubes on day 6 of differentiation. (c) Fusion index of myotubes. Images demonstrate myotubes (green: myosin heavy chain [MYH]) and nuclei (blue: 4’,6-diamidino-2-phenylindole [DAPI]). Myotubes are differentiated and cultured on 10–100 kPa gels and PS dishes. The fusion index is calculated as the percentage of nuclei in the myotubes relative to the total number of nuclei from the fluorescent staining image. Scale bars, 100 μm. Data are presented as the mean ± S.E.M., n = 25, \**p* < 0.05, vs. (> 1 GPa) (one-way analysis of variance (ANOVA) followed by Dunnett’s post hoc test).

### Effect of elasticity on differentiation of myoblasts

Subsequently, we assessed whether myoblasts seeded on flat gels can differentiate into myotubes. Myoblasts cultured on 10–100 kPa gels adhered and differentiated into myotubes, as those cultured on the PS dish for six days (Fig. 2b). In contrast, myotubes cultured on 160 and 230 kPa gels detached and formed cell aggregates by day 6 of differentiation (Sup. Fig. 2). Therefore, subsequent experiments were performed on 10–100 kPa gels. The fusion index from myoblasts to myotubes was calculated as the percentage of nuclei in myotubes relative to the total number of nuclei. The results demonstrated no difference in the fusion index for cells on 50 and 100 kPa gels compared to that on PS dishes, whereas cells on 10 and 30 kPa gels demonstrated a significantly lower index (Fig. 2c).

### Effect of elasticity on gene expression characterizing fiber types in myotubes

To assess the effect of culture substrate elasticity on the fiber type-characterizing gene expression in myotubes, we analyzed gene expression levels in myotubes cultured on 10–100 kPa gels and PS dishes. First, the gene expression levels of *MYH* isoforms (markers of fiber type) were assessed (Fig. 3a). The expression level of *MYH7*, a marker of type I fiber, was significantly increased in myotubes cultured on the gels of 10–50 kPa compared with that in the PS dish. The expression level of *MYH2*, a marker of type IIa fibers, exhibited no significant difference, although it tended to increase in myotubes cultured on gels with lower elasticity. In contrast, the expression levels of *MYH1* and *MYH4*, markers of type IIx and IIb fibers, respectively, did not differ among the PS dish and each gel with different elasticity. Subsequently, we assessed the expression levels of representative genes, markers of oxidative and glycolytic metabolism. Fig. 3b illustrates the expression levels of genes highly expressed in type I and IIa fibers that exhibit high oxidative metabolic capacity. The expression levels of *glucose transporter type 4* (*GLUT4*)—that transports glucose into the cell, and *myoglobin*—that supplies oxygen to mitochondria, were increased in myotubes cultured on the gels compared to those cultured on the PS dish. Specifically, myotubes cultured on 10 kPa gel exhibited the highest increase in gene expression. In contrast, *cytochrome c oxidase IV* (*COX IV*)—that is associated with ATP synthesis in mitochondria, exhibited no alteration in expression. We assessed *glyceraldehyde-3-phosphate dehydrogenase* (*GAPDH*), a glycolytic enzyme highly expressed in type IIx and IIb fibers that exhibit high glycolytic metabolic capacity. The expression level of *GAPDH* was upregulated in myotubes cultured on gels compared to that on PS dishes (Fig. 3c). *Peroxisome proliferator-activated receptor gamma coactivator 1-alpha* (*PGC-1α*) is a transcriptional coactivator that regulates the formation of type I and IIa fibers that exhibit high oxidative metabolic capacity. The expression level of *PGC-1α* was increased in myotubes cultured on the gels compared to that on the PS dish (Fig. 3d). Specifically, *PGC-1α* expression level was significantly increased in myotubes cultured on 10 kPa gel. Therefore, myotubes cultured on low-elasticity gels (10 kPa) exhibited enhanced expression of *PGC-1α*, myosin isoform (*MYH7* and *MYH2*), and oxidative metabolism-related genes (*GLUT4* and *myoglobin*), which characterize type I and type IIa fibers.

**Fig. 3.**
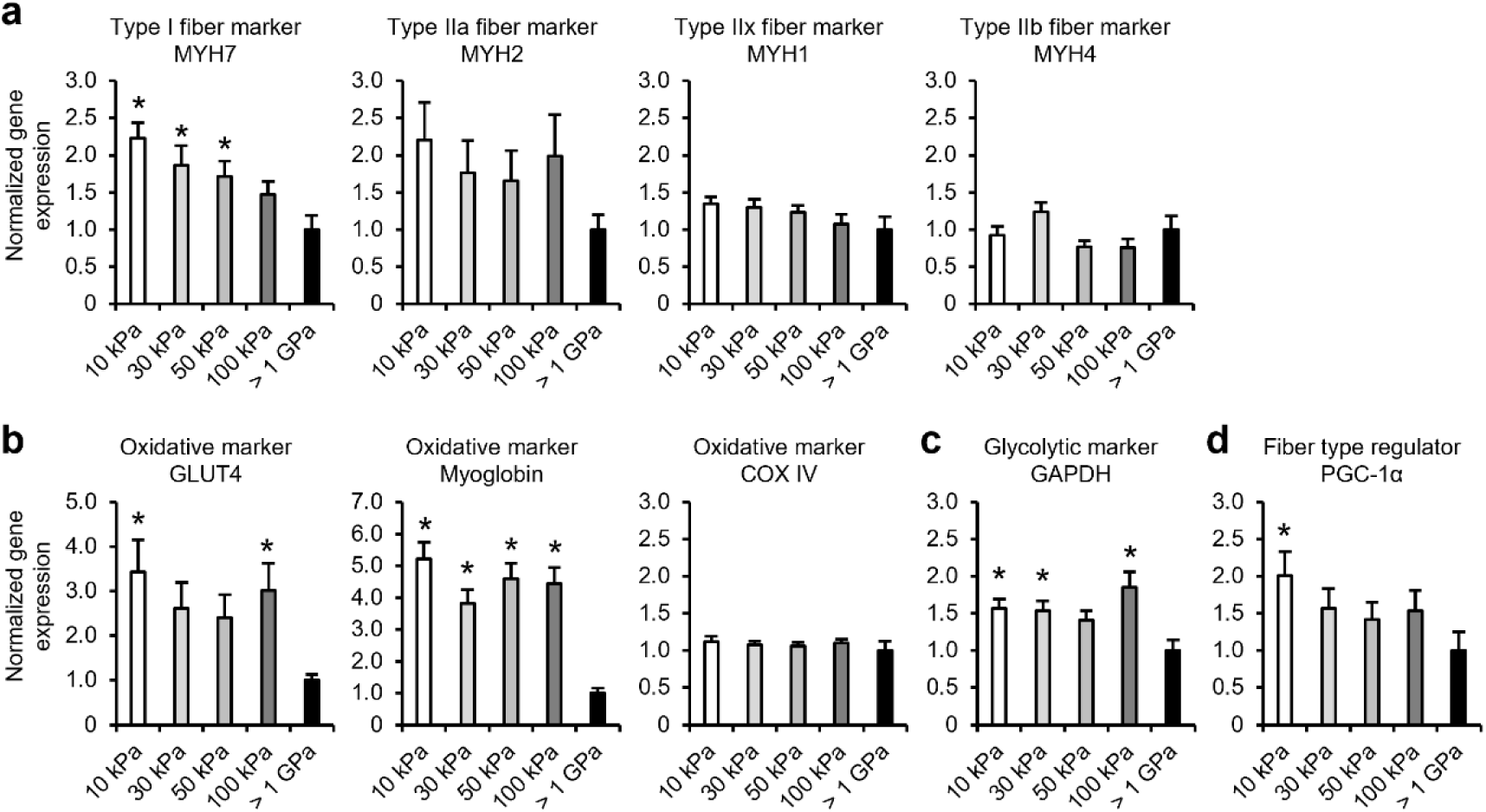
Gene expression of myotubes cultured on flat gels with different elasticity. Gene expression levels of myotubes differentiated and cultured on 10–100 kPa gels with a flat surface and PS dish (> 1 GPa) for six days are analyzed using qRT-PCR. (a) Expression levels of *MYH* isoform genes in myotubes. The graphs demonstrate the expression levels of *MYH7*, *MYH 2*, *MYH 1*, and *MYH 4*, which are marker genes for fiber types. (b) Expression levels of genes related to oxidative metabolism. The graphs demonstrate the expression levels of *glucose transporter type 4* (*GLUT4*), *myoglobin*, and *cytochrome c oxidase IV* (*COX IV*), which are marker genes for oxidative metabolism. (c) Expression levels of genes related to glycolytic metabolism. The graph demonstrates the expression level of *glyceraldehyde-3-phosphate dehydrogenase* (*GAPDH*), which is a marker gene for glycolytic metabolism. (d) Expression levels of *peroxisome proliferator-activated receptor gamma coactivator 1-alpha* (*PGC-1α*) gene, which regulates the formation of type I and IIa fibers, which exhibit high oxidative metabolic capacity. The graph demonstrates the expression level of *PGC-1α*. Expression levels of each gene are standardized using the housekeeping gene *hypoxanthine phosphoribosyltransferase 1* (*Hrpt1*). Data are presented as the mean ± S.E.M., n = 5–6, \**p* < 0.05, vs. (> 1 GPa) (one-way ANOVA followed by Dunnett’s post hoc test).

### Effect of microgrooves on myoblast differentiation

Microgrooved materials, such as poly (lactic-co-glycolic acid) and gelatin methacrylate facilitate orientation and *MYH* expression [22, 26]. Therefore, we comprehensively analyzed whether myoblasts differentiate on gels with elasticities of 10, 30, 50, or 100 kPa fabricated with microgroove widths of 3, 5, 10, or 50 μm. Myoblasts differentiated into myotubes on each microgroove, similar to the flat condition, regardless of elasticity (Fig. 4a). The fusion index was significantly increased in 3–10 μm width grooves compared to that in the flat condition, regardless of gel elasticity (Fig. 4b). Grooves with a width of 50 μm significantly increased the fusion index of myotubes only in 50 kPa gel. The orientation of differentiated myotubes along the grooves increased in each groove width compared to that under the flat condition, regardless of gel elasticity (Fig. 4c).

**Fig. 4.**
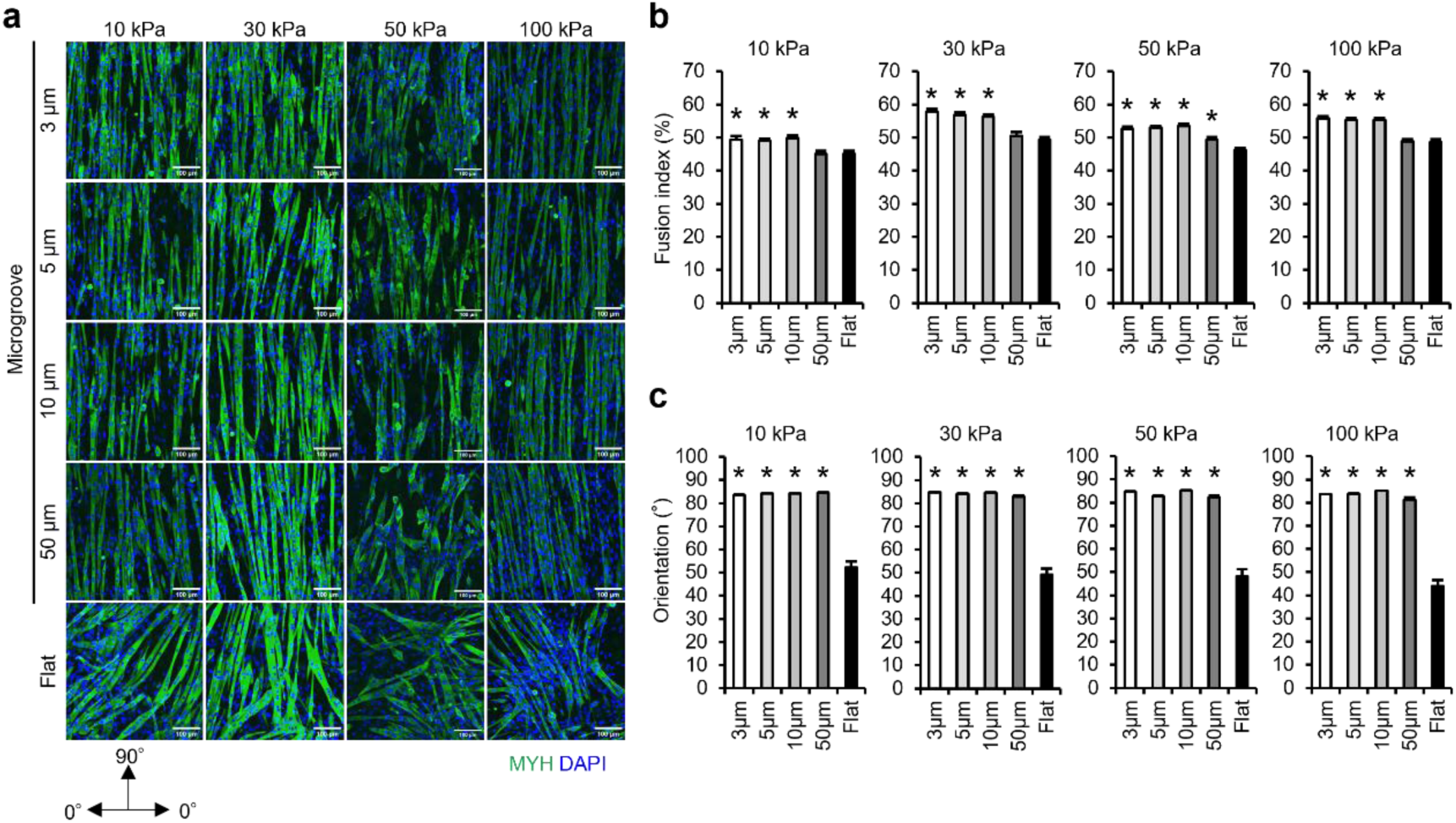
Fusion index and orientation of myotubes cultured on gelatin gels with transferred microgrooves. (a) Fluorescent images of myotubes differentiated and cultured on 10–100 kPa gelatin gels with transferred 3, 5, 10, and 50 μm microgrooves and flat condition for six days. The images demonstrate differentiated myotubes (green: MYH) and nuclei (blue: DAPI). (b) Fusion index of myotubes. The fusion index is calculated as the percentage of nuclei (blue) in the myotubes (green) relative to the total number of nuclei. (c) Orientation of myotubes. Orientation is analyzed from the longitudinal axis direction of myotubes, with the vertical direction of the microgroove as 0° and the parallel direction as 90°. Scale bars, 100 μm. Data are presented as the mean ± S.E.M., n = 45–100 (images), \**p* < 0.05, vs. (Flat) (one-way ANOVA followed by Dunnett’s post hoc test).

### Effect of microgrooves on gene expression characterizing fiber types in myotubes

To assess the effect of microgrooves on fiber type-characterizing gene expression in myotubes, we analyzed gene expression levels in myotubes cultured on gels with groove widths ranging from 3–50 μm. First, the gene expression levels of *MYH* isoforms were assessed. The expression levels of *MYH7* and *MYH1* were significantly increased or tended to increase in the 3 μm groove compared with that in the flat condition, regardless of gel elasticity (Fig. 5a). Additionally, there was a reduced *MYH2* expression at 50 μm groove, specifically in the 50 kPa gel—that was significantly reduced. *MYH4* expression level was unaltered by grooves. Subsequently, we assessed the expression levels of genes associated with oxidative and glycolytic metabolism. *GLUT4* expression levels remained unaltered (Sup. Fig. 3a). *Myoglobin* expression levels tended to increase and decrease in the 3 and 50 μm grooves, respectively, compared with that in the flat condition, with no significant difference. *COX IV* expression level tended to decrease with groove condition and was significantly decreased in the 5–10 µm grooves only in 100 kPa gels compared with that in the flat condition. *GAPDH* expression level was unaltered (Sup. Fig. 3b). *PGC-1α* expression reduced as the groove width increased, such as 50 μm, regardless of gel elasticity; however, there was no significant difference (Fig. 5b). Therefore, gels on which microgrooves were transferred, regardless of elasticity, altered the expression levels of certain *MYH* isoforms and metabolism-related genes in myotubes but did not induce a shift to the genetic characteristics of each fiber type.

**Fig. 5.**
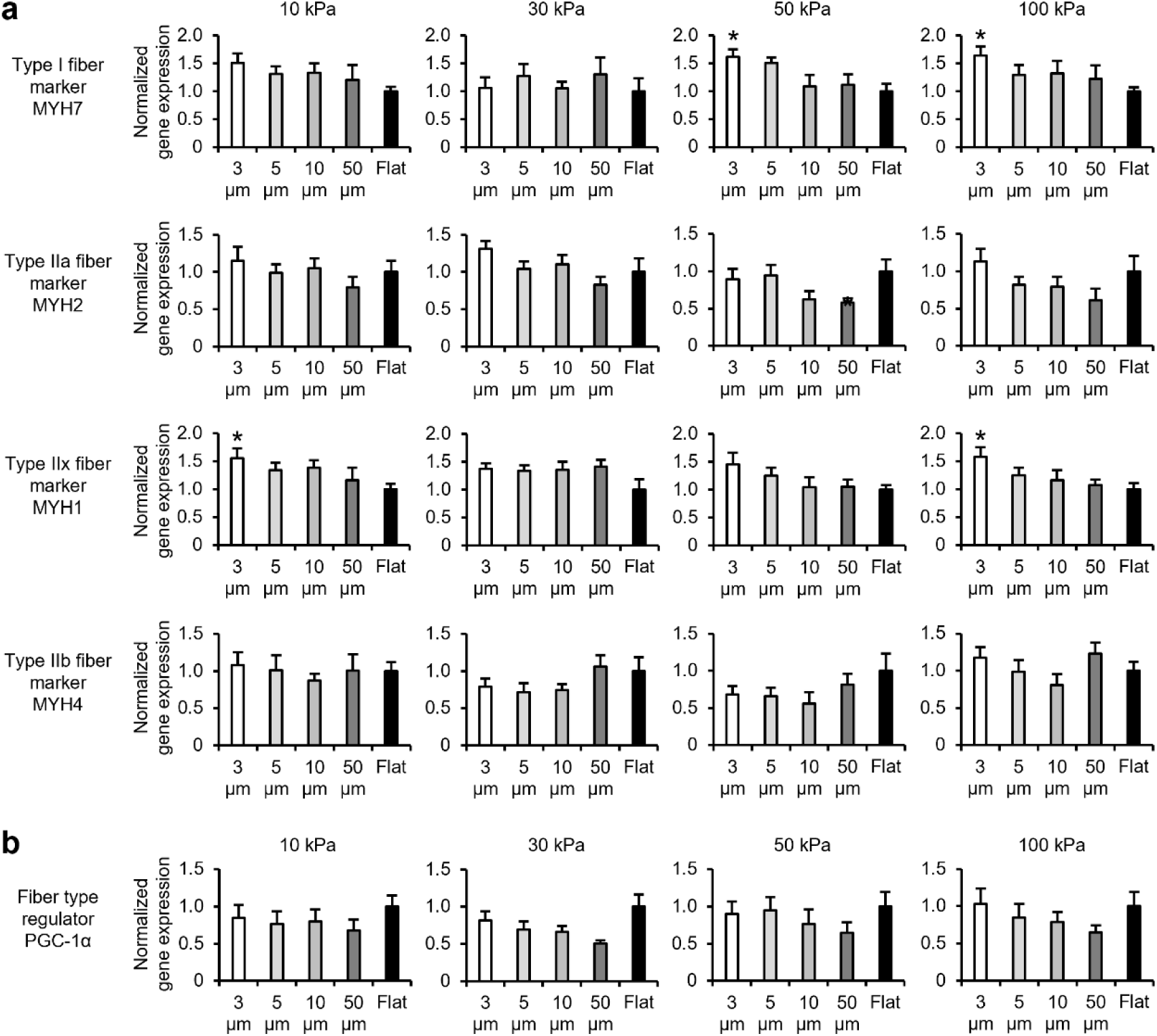
Gene expression of myotubes cultured on gelatin gels with transferred microgrooves. Gene expression in myotubes differentiated and cultured on 10–100 kPa gels with transferred 3, 5, 10, and 50 μm microgrooves and flat conditions for six days is analyzed using qRT-PCR. (a) Expression levels of *MYH* isoform genes in myotubes. The graphs demonstrate the expression levels of *MYH7*, *MYH 2*, *MYH 1*, and *MYH 4*, which are marker genes for fiber types. (b) Expression levels of *PGC-1α* gene, which regulates the formation of type I and IIa fibers that exhibit high oxidative metabolic capacity. The graphs demonstrate the expression level of *PGC-1α*. Expression levels of each gene are standardized using the housekeeping gene *Hrpt1*. Data are presented as the mean ± S.E.M., n = 7–8, \**p* < 0.05, vs. (Flat) (one-way ANOVA followed by Dunnett’s post hoc test).

## Discussion

It has been observed that the elasticity and microstructure of the culture substrate, mimicking the *in vivo* cell niche, enhance myotube development, differentiation, and orientation. Consequently, extensive research has been conducted to enhance these effects and elucidate the mechanisms using various substrates. However, to the best of our knowledge, no studies assessed whether these mechanical stimuli induce a shift of myotubes to the characteristics of each fiber type. In this study, we used a crosslinked gelatin gel prepared using a radiation-induced reaction to comprehensively assess the effects of elasticity and microgrooves. Our findings, for the first time, provided evidence that (i) an elasticity of 10 kPa induces myotube shifts toward the genetic characteristics of type I and IIa fibers that exhibit high oxidative metabolic capacity, and (ii) microgrooves with 3–10 µm width enhance myoblast fusion and myotube orientation compared to those with larger widths, without inducing the shift to the genetic characteristics of each fiber type.

Previously, artificial polymers, such as PDMS and polyacrylamide were used to prepare culture substrates with tunable elasticity and microgrooves. However, these materials should be coated with ECM extract reagents to facilitate cell adhesion on the surface. The concentration and distribution of bioactive ligands differ depending on the coating method and type of substrate material, complicating the assessment of mechanical stimuli [37, 38]. When using ECM-derived proteins as the base material for culture substrates, the processing method becomes a concern. Glutaraldehyde and carbodiimide are commonly used to crosslink proteins, but these reagents consume the cell-binding motifs of ECM-derived proteins during crosslinking [39, 40]. Although a microbial transglutaminase-crosslinked gel was developed [25] as an alternative, its tunable elasticity range is limited [41]. In contrast, our crosslinking method using OH radicals—a product of water radiolysis—does not require any chemical reagent. In this method, tyrosine, phenylalanine, and histidine serve as crosslinking sites for gelatin, thereby preserving RGD (arginine, glutamic acid, and aspartic acid) cell-binding motifs of gelatin even after processing [36, 42]. Additionally, the elasticity and micropatterns can be controlled over a wide range, facilitating a comprehensive analysis of how mechanical stimuli influence the shift of myotubes to the characteristics of each fiber type.

To assess the effect of culture substrate elasticity on the induction of myotube shifts to each fiber type, the substrate should have high cell adhesion and cytocompatibility. Myoblasts adhered to the crosslinked gelatin gels regardless of the elastic modulus, and over 99 % survived for 48 h post-seeding (Fig. 2a). Upon differentiation, myoblasts fused into myotubes on gels with an elastic modulus of 10–100 kPa and on PS dishes with a modulus of ≥ 1 GPa (Fig. 2b). In contrast, on gels with 160 and 230 kPa, myoblasts adhered to the gel at 24 h post-seeding, but consistently detached and formed aggregates by day 6 of differentiation (Sup. Fig. 2). The detachment and formation of aggregates are explained by the cell-substrate and cellular traction forces [43]. The cell-substrate force is enhanced by the formation of stress fibers composed of non-muscle myosin and actin filaments linked to focal adhesions, based on the increase in elasticity of substrates [44]. Additionally, the traction force is enhanced by the formation of the total amount of intracellular stress fibers based on the increase in elasticity of substrates [45]. Therefore, on stiff substrates, where the increased traction force exceeds the cell-substrate adhesion force, cells use the traction force to pull themselves away from the substrate and aggregate [43, 46]. In contrast, gels with an elasticity of ≥ 160 kPa are excessively stiff for culturing myotubes owing to a lower cell-substrate adhesion force than that of the traction force. Gels with an elasticity of ≤ 100 kPa are optimal for culturing myotubes without detachment because the cell-substrate adhesion force is dominant. Notably, this aligns with the fact that an elastic modulus of 10–100 kPa is comparable to the range of elasticity of skeletal muscle [31–35]. Additionally, high-stiffness PS dishes with collagen coating to enhance cell adhesion have higher cell-substrate adhesion force than that of traction force; therefore, cells can be cultured on a flat surface without aggregation [43].

The fiber type is determined by the expression levels of *MYH* isoforms that are crucial for muscle contraction. Type I, IIa, IIx, and IIb fibers have high expression levels of *MYH7*, *MYH2*, *MYH1*, and *MYH4*, respectively [3]. In this study, as the elasticity of the gelatin gel reduced from 100 to 10 kPa, the gene expression levels of *MYH7* and *MYH2* increased in myotubes cultured on the gel (Fig. 3a). Therefore, the elasticity demonstrated a shift of myotubes to the genetic characteristic of type I and IIa fibers. Additionally, type I and IIa fibers are associated with glucose tolerance and oxidative metabolism [3]. Slow-twitch muscles—primarily composed of type I and IIa fibers—have a high expression of metabolic-related genes, such as *GLUT4* (transports glucose into the cell), *myoglobin* (oxygen supply to mitochondria), and *COX IV* (oxidative phosphorylation in the mitochondria) [47, 48]. Myotubes cultured on gelatin gels increased the expression of *myoglobin* and *GLUT4* (Fig. 3b). This indicates that reduced gel elasticity induces myotube shifts to the genetic characteristics of type I and IIa fibers, altering both *MYH* isoforms and metabolic characteristic-associated gene expression. The shift to each fiber type is controlled by gene expression through transcription factors. For example, *PGC-1α*—a transcription coactivator—regulates the expression of genes involved in oxidative metabolism and induces myofibers to become type I and IIa fibers [49, 50]. In L6 rat myoblast-derived myotubes, *PGC-1α* overexpression enhances the expression of *GLUT4* and *COX IV* that are highly expressed in type I and IIa fibers with high oxidative metabolic capacity [50]. Additionally, mice with physiologically overexpressing *PGC-1α* levels in skeletal muscles demonstrated increased expression of *myoglobin* and *COX IV* compared to that of the wild-type mice, and increased presence of type I and IIa fibers in the skeletal muscle [49]. This study demonstrated that *PGC-1α* expression in myotubes increased in gelatin gels with a low modulus of elasticity, such as 10 kPa (Fig. 3d). The mRNA expression level of *PGC-1α* is suppressed by activated Yes-associated protein (YAP) [51]. YAP—a transcriptional coactivator—is less active when localized in the cytoplasm and more active when localized in the nuclei. The subcellular localization of YAP is regulated by culture substrate elasticity—the cells cultured on a softer culture substrate localize to the cytoplasm and have reduced activity [52]. This indicates that myotubes cultured in a lower-elasticity gelatin gel exhibit increased *PGC-1α* expression owing to reduced YAP activity, leading to a shift toward the genetic characteristics of type I and IIa fibers. Notably, in this study, *COX IV* expression, which is highly expressed in type I and IIa fibers, did not depend on gel elasticity (Fig. 3b). Because *COX IV* expression is increased by *PGC-1α* [48], it is indicated that myotubes cultured on gels have regulatory factors that differ from *PGC-1α*. For example, cells cultured on soft gels (0.5 kPa polyacrylamide) demonstrate increased nuclear localization and activation of the transcriptional co-activator *histone deacetylase 4* (*HDAC4*) compared to those cultured on stiff gels (100 kPa) [53]. Because activated *HDAC4* suppresses *COX IV* expression, it was indicated that substrate elasticity has a mechanism that controls the expression of metabolic-related genes through multiple signaling pathways [54]. Further studies are required to assess the mechanism of elasticity-induced shift of myotubes to the genetic characteristics of type I and IIa fibers.

Slow- and fast-twitch muscles differ in their elasticity. In mice, the elasticity of myofibers isolated from soleus muscles (slow-twitch muscles) is greater than that isolated from extensor digitorum longus (fast-twitch muscles) [35]. Additionally, ECM elasticity is affected by collagen I content and crosslinking density [55]. Slow-twitch muscles generally contain more collagen I than that of fast-twitch muscles. The crosslinking density of collagen I is higher, indicating that the ECM in slow-twitch muscles is stiffer than that in fast-twitch muscles [56]. Therefore, if the elasticity of slow- and fast-twitch muscles facilitates the shift of myotubes to the genetic characteristics of each fiber type, the substrate with higher elasticity may induce a shift of myotubes to type I and IIa fibers, characterizing slow-twitch muscles. However, in this study, the gelatin gel with the lowest elasticity (10 kPa) within the range of elasticity in skeletal muscle (10–100 kPa) [31–35] induced a shift of myotubes to the genetic characteristics of type I and IIa fibers. This indicates that the differences in the composition of fiber types in each skeletal muscle are not induced by the elasticity of mature muscle tissues. In contrast, the results explain the shift of skeletal muscle cells to each fiber type during development. During early development, skeletal muscles contain a large amount of collagen III that contributes less to elasticity than that of collagen I. As muscles mature, collagen III shift to collagen I [57]. Therefore, the elasticity derived from collagen in skeletal muscles during development may be lower than that in mature skeletal muscles. During skeletal muscle development, type I fibers are formed first, followed by type IIa, IIx, and IIb fibers [3, 4]. The relationship between elasticity and the induction of type I fibers during development aligns with the mechanosensing mechanism of myotubes identified in this study.

Microgrooves on culture substrates enhance myotube differentiation and orientation [22]. In this study, a comprehensive analysis demonstrated that microgrooves (3–10 μm) enhanced the fusion index and orientation of myotubes compared with that of the flat condition, regardless of elasticity (Fig. 4b and c). This effect likely stems from the action of microgrooves on myoblasts before differentiation. In 10–100 kPa gels, myoblasts before differentiation elongated parallel to the microgrooves (3–10 μm) and exhibited a high aspect ratio and longitudinal length compared with those under the flat condition (Sup. Fig. 4a-d)—similar to non-muscle cells (HeLa, Madin-Darby Canine Kidney, and 3T3-Swiss cells) observed in our previous study [36]. When myoblasts come into contact with each other, they fuse and differentiate into myotubes [58]. Myoblasts aligned along the groove do not fuse when their sides come into contact with each other but initiate fusion when their elongated ends meet [59]. Therefore, it is indicated that microgrooves (3–10 μm) enhanced their fusion index and orientation by increasing the opportunities for the ends of myoblasts elongated in the line direction to come into contact with each other.

The effect of microgrooves on culture substrates in inducing a shift of myotubes to the characteristics of each fiber type has not been confirmed. Myotubes cultured on microgrooves with 3–10 μm width exhibited a significant increase in the gene expression of *MYH7*, *MYH2*, and *MYH1* compared with that in flat conditions, or a trend towards such an increase (Fig. 5a). Additionally, there were no significant alterations in the expression levels of *myoglobin*, *GLUT4*, *COX IV*, or *GAPDH*—that are associated with metabolism (Sup. Fig. 3a and b). Therefore, we concluded that microgroove-derived mechanical stimulation enhances the fusion index and orientation of myotubes and increases the expression level of *MYH* isoforms but does not contribute to the shift of myotubes to the genetic characteristics of each fiber type. One reason the microgrooves did not induce the shift of myotubes to the genetic characteristics of each fiber type is that they localize YAP to the nuclei [60]. When YAP is localized to the nucleus, it is activated, and *PGC-1α* expression is suppressed. In our results, the amount of *PGC-1α* expression tended to decrease as the groove width increased (Fig. 5b), indicating that microgrooves did not alter the expression of metabolic-related genes that characterize fiber types because they failed to increase *PGC-1α* expression.

In this study, we comprehensively analyzed the combined effects of elasticity and microgrooves on the induction of shifts to each fiber type using gelatin gels prepared with radiation-induced reactions. Myotubes exhibited enhanced expression of *PGC-1α*, *MYH* isoforms, and metabolic-related genes characteristic of type I and IIa fibers that exhibit oxidative metabolism at low gel elasticity (10 kPa) (Fig. 6a-c). In contrast, although the expression of *MYH* isoforms was enhanced by 3 µm microgrooves, the expression of metabolic-related genes and *PGC-1α* did not increase (Fig. 6a-c). Additionally, it was observed that microgrooves did not inhibit the induction of elasticity-induced myotube shifts to the genetic characteristics of type I and IIa fibers. Therefore, we concluded that elasticity induces myotube shifts toward the genetic characteristics of oxidative type I and IIa fibers, and microgrooves enhance fusion index and orientation (Fig. 6d). It was assumed that myofiber type in skeletal muscle cells is primarily determined by body parts and genetics, leading to shifts between distinct fiber types. Moreover, fiber types shift due to disease and chronic exercise. This study demonstrates that substrate elasticity mimicking cell niches induces muscle fiber-type shifts. Further studies should elucidate the mechanism by which culture substrate elasticity induces the shift of myotubes to the genetic characteristics of type I and IIa fibers. This study demonstrated a method to create highly oriented slow-twitch muscle models composed of type I and IIa fibers that exhibit oxidative metabolism by controlling the elasticity and microgrooves of gelatin gel. These models can help elucidate the mechanism by which slow-twitch muscles preferentially lose muscle mass and strength in bedridden patients or those with cancer. Furthermore, they can be used for drug screening for prevention and treatment. Aligned myotubes exhibit effective therapeutic effects after muscle transplantation [61, 62]. Because of the biocompatibility and biodegradability of our crosslinked gelatin gels [36], it is anticipated that artificial slow-twitch muscle tissues on the gels can be transplanted into the body and may be useful as transplanted therapeutic materials for severe muscle damage, specifically slow-twitch muscle damage.

**Fig. 6.**
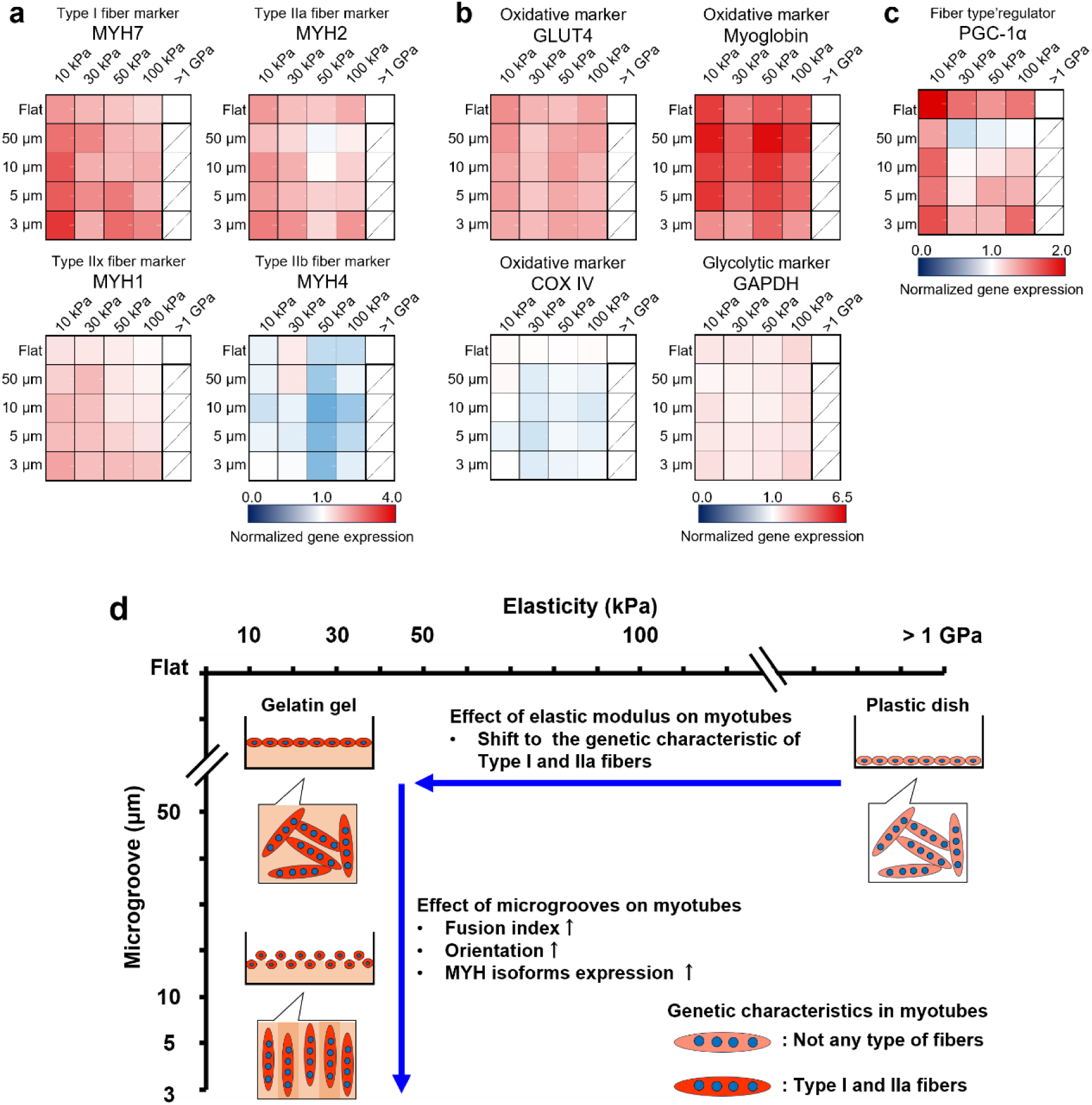
Summary of the effects of elasticity and the microgrooves of the culture substrate on the induction of myotube shifts to each fiber type. (a) Heatmaps of gene expression levels of *MYH* isoforms, which are markers of fiber types. (b) Heatmaps of gene expression levels related to oxidative and glycolytic metabolism. (c) Heatmap of gene expression levels of *PGC-1α*, which regulates the formation of type I and IIa fibers that exhibit high oxidative metabolic capacity. These gene expressions in myotubes differentiated and cultured on plastic (PS) dishes (> 1 GPa) and 10–100 kPa gels with 3, 5, 10, and 50 μm microgrooves and flat conditions for six days are analyzed using qRT-PCR. The expression levels of each gene are standardized using the housekeeping gene *Hrpt1*. The heatmaps are normalized by the gene expression levels of myotubes cultured on the PS dish. Data are presented as the mean ± S.E.M., n = 5–8. (d) The gelatin gel with low elasticity, such as 10 kPa, induces the shift of myotubes to the genetic characteristic of type I and IIa fibers which exhibit high oxidative metabolic capacity. The microgrooves, such as 3 μm width, transferred to the gelatin gel surface increased the fusion index, orientation, and expression level of *MYH* isoforms in myotubes. The effects of elasticity and microgrooves of culture substrates are independent, and by creating a gelatin gel with a combination of any elasticity and microgrooves, the myotubes that exhibit the characteristics desired by the researchers can be cultured.

## Materials and Methods

### Preparation of gelatin gel using radiation

The preparation of crosslinked gelatin gels using γ-ray irradiation-induced chemical reactions is illustrated in Fig. 1a. Gelatin (porcine skin, Type A. GLS250; Nitta Gelatin, Osaka, Japan) solution was prepared by dissolving 10 wt% (for 8 and 10 kGy irradiation) or 15 wt% (for 15, 20, 25, and 30 kGy irradiation) in ultrapure water and heating at 50 °C for 30 min. The solution was poured into 24 well plates or 35 mm dishes, sealed in a plastic bag with a deoxygenant (A-500HS; AS ONE, Osaka, Japan), and incubated at 20 °C overnight for physical gelation. To form microgrooves on the gel surface, a PDMS mold was placed on the gelatin solution before physical gelation. The PDMS mold with 2 μm depth and 3, 5, 10, or 50 μm width microgrooves was prepared as follows. The base polymer of PDMS (SIM-260) and crosslinker (CAT-260) (both from Shin-Etsu Chemical, Tokyo, Japan) were mixed in a 10:1 ratio, applied evenly to a silicon master mold (DTM-1-1; Kyodo International, Kanagawa, Japan), and degassed. The PDMS mold was cured at 150 °C for 30 min and subsequently peeled off from the master mold. The physical gelatin gel was irradiated with γ-rays supplied by a ^60^Co No. 2 Irradiation Facility at the Takasaki Institute for Advanced Quantum Science, QST, at a dose rate of 4 or 5 kGy/h at 15–20 °C. This process transformed the physical gelatin gel into a sterilized and crosslinked gelatin gel without chemical treatment. After the PDMS mold was removed, the irradiated gel was washed twice with phosphate-buffered saline (PBS) (-), first for 1 h at 37 °C and then overnight (Fig. 1a). The PBS-washed gels were cultured in a growth medium at 37 °C for two 1-hour cycles to remove residual PBS before cell culture. The elasticity of the flat gelatin gels was measured using a rheometer (RE2-33005C; Yamaden, Tokyo, Japan) after washing twice with PBS. A cylinder with a diameter of 3 mm was pressed against the gel surface at a speed of 50 μm/s using a 2 N load cell, and the slope of the approximate straight line of the stress-strain curve was calculated as the compressive elasticity. Microgrooves on the surface of the gelatin gel were imaged using an inverted microscope (IX83; Evident, Tokyo, Japan).

### Cell culture

The C2C12 myoblast cell line (EC91031101; DS Pharma Biomedical, Osaka, Japan) was seeded onto dishes coated with Cellmatrix (Type I-A, Nitta Gelatin) or gelatin gels and cultured in a growth medium consisting of Dulbecco’s modified Eagle’s medium (DMEM) (high-glucose DMEM, 08488-55; Nacalai Tesque, Kyoto, Japan) supplemented with 10 % fetal bovine serum (12483020; Thermo Fisher Scientific, MA, U.S.A.), 2 mM L-Glutamin (G7513; Merck, Darmstadt, Germany), 100 U/mL penicillin, and 100 µg/mL streptomycin (15140122; Thermo Fisher Scientific) at 37 °C with 5 % CO_2_. After one day, the medium was replaced with a differentiation medium consisting of DMEM supplemented with 2 % calf serum (16010167; Thermo Fisher Scientific), 2 mM L-glutamine, minimum essential medium non-essential amino acids solution (139-15651; FUJIFILM Wako Chemicals, Osaka, Japan), and 100 U/mL penicillin. Half of the medium was replaced daily.

### Analysis of myoblast viability, proliferative rate, area, orientation, aspect ratio, and longitudinal length

C2C12 myoblasts were seeded on 24-well plates and gelatin gels at a density of 5.3 × 10^3^ cells/cm^2^ and cultured in a growth medium for 24 or 48 h. To assess the viability and proliferative rate of myoblasts, live cells were stained with calcein-AM (C396; Dojindo Laboratories, Kumamoto, Japan), dead cells were stained with propidium iodide (P378; Dojindo Laboratories), and nuclei were stained with Hoechst 33342 (H342; Dojindo Laboratories). Myoblast viability at 48 h was calculated as the percentage of live cells among the total number of live and dead cells using a fluorescence microscope (BZ-X800; Keyence, Osaka, Japan). The proliferation rate of myoblasts per 24 h was calculated by dividing the number of nuclei at 24 h by the number of nuclei at 48 h of culture. To assess the myoblast area, myoblasts were stained with Acti-stain 555 phalloidin (PHDH1-A; Dojindo Laboratories) 24 h post-seeding. Fluorescent images of the stained cells were obtained using an upright microscope (BX51WI+BX-DSU; Evident) connected to a 10x water immersion objective (UMPLFLN10XW; Evident) and a digital camera (ORCA-Flash4.0 V3; Hamamatsu Photonics, Shizuoka, Japan). Myoblast area, orientation (angle from 0–90◦), aspect ratio, and longitudinal length were analyzed based on myoblasts stained with phalloidin using ImageJ-Fiji.

### Assessment of myotube fusion index and orientation

C2C12 myoblasts were seeded at a density of 4.2 × 10^4^ cells/cm^2^ on 35 mm dishes and gelatin gels and cultured for six days. Cultured myotubes were fixed with 4 % paraformaldehyde (163-20145; FUJIFILM Wako) for 10 min and incubated with 0.3 % Triton X-100 (35501-02; Nacalai Tesque) and 1 % bovine serum albumin (P6154-100G; Biowest, Nuaillé, France). Nuaillé, FRANCE) in PBS for 30 min at room temperature. Subsequently, myotubes were incubated with MYH antibody (1:300, 14-6503-82; Thermo Fisher Scientific) overnight at 4 °C and labeled with Alexa Fluor 488 (1:300, A28175; Thermo Fisher Scientific) for 1 h at room temperature. Nuclei were labeled with 4’,6-diamidino-2-phenylindole (DAPI, 340-07971; Dojindo Laboratories). Fluorescent images of stained cells were obtained from random fields of view for each substrate using an upright microscope. The fusion index was calculated as the percentage of nuclei in myotubes relative to the total number of nuclei, using ImageJ-Fiji. The parameters for calculating the nuclei were modified from those described previously [63]. The orientation of myotubes (angle from 0–90◦) was calculated using the Directionality plugin in ImageJ-Fiji.

### mRNA extraction and quantitative real-time polymerase chain reaction (qRT-PCR)

C2C12 myoblasts were seeded at a density of 4.2×10^4^ cells/cm^2^ on 35 mm dishes or gelatin gels and differentiated for six days. mRNA was extracted from myotubes using NucleoSpin🄬 RNA Plus XS (U0990B; Takara Bio, Shiga, Japan). mRNA was reverse transcribed to cDNA using PrimeScript™ RT Master Mix (RR036A; Takara Bio) in a thermal cycler (TP350; Takara Bio). The qRT-PCR was performed using a real-time qPCR system (CFX96; Bio-Rad) and TB Green🄬 Fast qPCR Mix (RR430A; Takara Bio). Primer sequences (synthesized by Eurofins Genomics, Tokyo, Japan) for this study are listed in Sup. Tab.1. mRNA expression levels of each gene were normalized to the housekeeping gene *hypoxanthine phosphoribosyltransferase 1* (*Hprt1*) and determined using the cycle threshold values with the ⊿⊿Ct method.

### Statistics

Data are presented as the mean ± standard error of the mean. For multiple comparisons, data were analyzed using one-way analysis of variance, followed by Dunnett’s post hoc test. Statistical significance was set at *p* < 0.05.

## Data availability statement

All data that supports the findings of this study are included within the article and any supplementary file and movies.

## Acknowledgments

The authors thank Ms. Ryoko Mezaki (QST), Ms. Yukiko Sakatsume (QST), and Ms. Noriko Tawara (QST) for technical assistance. This research was supported by the Innovative Science and Technology Initiative for Security, Acquisition, Technology, and Logistics Agency (Grant No. JPJ004596, ATLA, Japan), JSPS KAKENHI (Grant Nos. 22H05054, 23K19377, and 24K01998) and ACT-X (Grant No. JPMJAX2014), and A-STEP (Grant No. JPMJTR22U7) from the Japan Science and Technology Agency (JST). We would like to thank Editage for English language editing.

## Contributions

H.H., T.G.O., K.O., Y.M., N.L.F., and M.T. designed the research. H.H. and T.G.O. performed the experiments and analyzed the data. All authors wrote the manuscript and approved the final manuscript.

