## Supplementary Information for "Combined stimuli of elasticity and microgrooves form aligned myotubes that characterize slow-twitch muscles"

**Supplementary Figures**

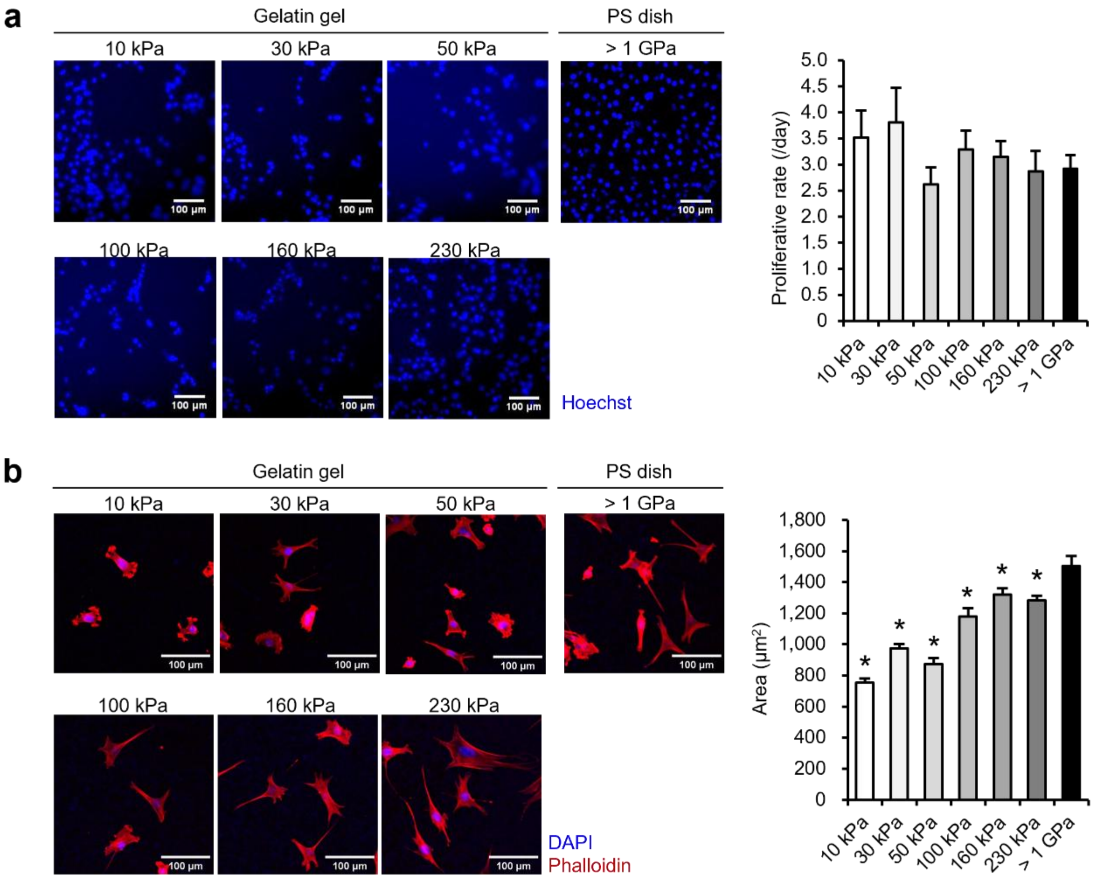


**Sup. Fig. 1 Proliferative rate and area of myoblasts seeded on flat gelatin gels.**

(a) Proliferative rate of myoblasts cultured on 10–230 kPa gels and plastic dishes (PS dish, > 1 GPa). The images demonstrate myoblasts at 48 h post-seeding stained with nuclei (blue, Hoechst 33342). The proliferative rate is calculated by dividing the number of cells at 48 h post-seeding by the number of cells at 24 h. (b) Area of myoblasts cultured on 10–230 kPa gels and PS dishes. The images demonstrate myoblasts at 24 h post-seeding, stained for actin (red: phalloidin) and nuclei (blue: 4',6-diamidino-2-phenylindole (DAPI)). The area is calculated as the cell area surrounded by stained actin. Scale bars, 100 µm. Data are presented as the mean ± standard error of the mean (S.E.M.), n = 10 (a) or 25 (b), *$p < 0.05$, vs. (> 1 GPa) (one-way analysis of variance (ANOVA) followed by Dunnett's post hoc test).

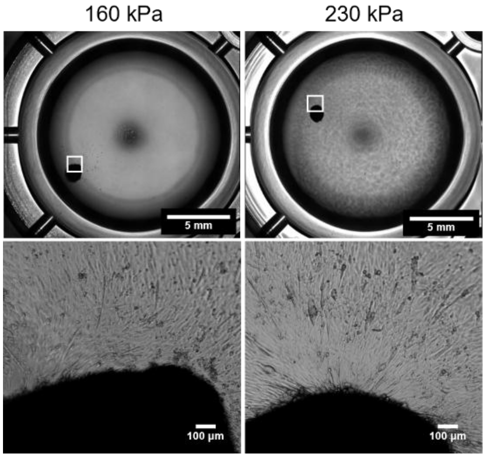


**Sup. Fig. 2 Myotubes differentiated on 160 and 230 kPa gelatin gels.**

Phase contrast images of myotubes differentiated on 160 and 230 kPa gelatin gels with flat surfaces for six days. The upper images demonstrate the whole well. Scale bars, 5 mm. The lower images demonstrate an enlarged cell aggregate of the square area. Scale bars, 100 µm.

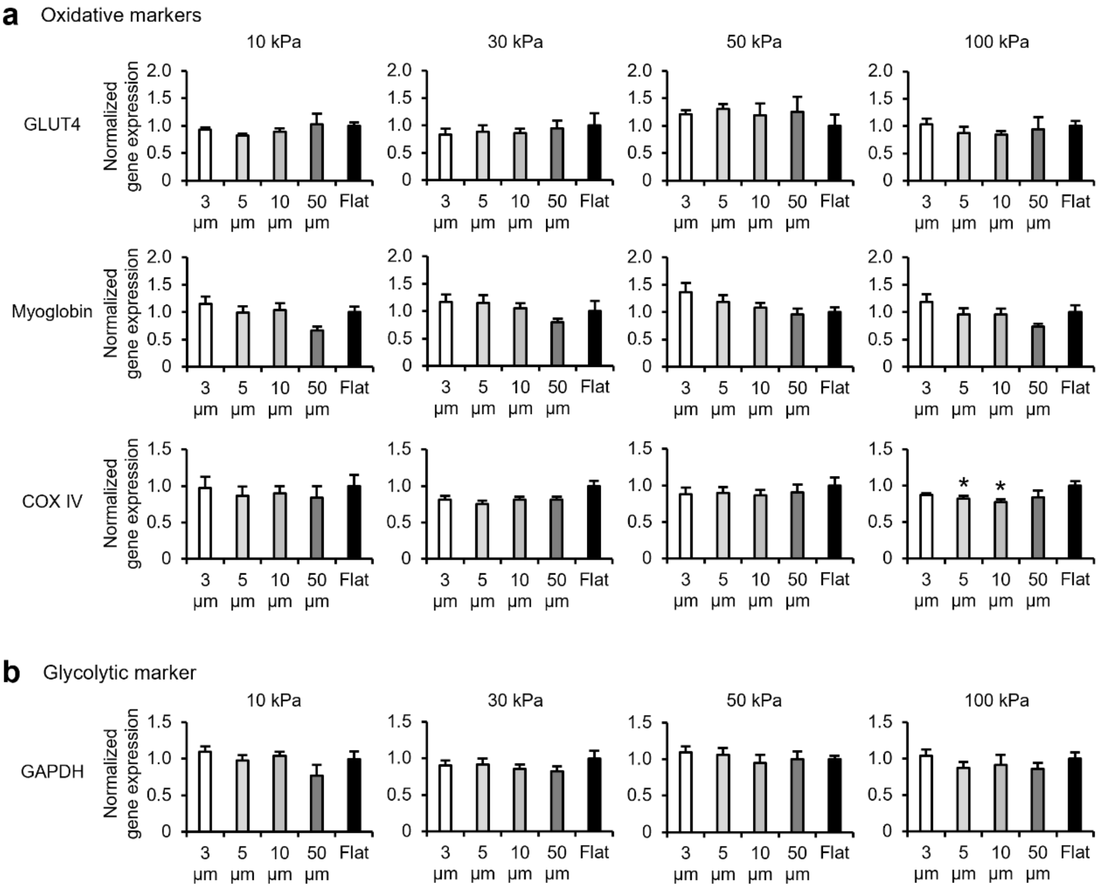


**Sup. Fig. 3 Metabolic-related gene expression of myotubes cultured on gelatin gels with transferred microgrooves.**

Metabolic-related gene expression of myotubes differentiated on 10–100 kPa gels with 3, 5, 10, and 50 µm microgrooves and flat condition for six days is analyzed using qRT-PCR. (a) Expression level of genes related to oxidative metabolism. The graphs demonstrate the expression levels of *glucose transporter type 4* (*GLUT4*), *myoglobin*, and *cytochrome c oxidase IV* (*COX IV*), which are marker genes for oxidative metabolism. (b) Expression level of genes related to glycolytic metabolism. The graphs demonstrate the expression level of *glyceraldehyde-3-phosphate dehydrogenase* (*GAPDH*), which is a marker gene for glycolytic metabolism. Expression levels of each gene are standardized using the housekeeping gene *hypoxanthine phosphoribosyltransferase 1* (*Hrpt1*). Data are presented as the mean ± S.E.M, n = 7–8, *$p < 0.05$, vs. (Flat) (one-way ANOVA followed by Dunnett's post hoc test).

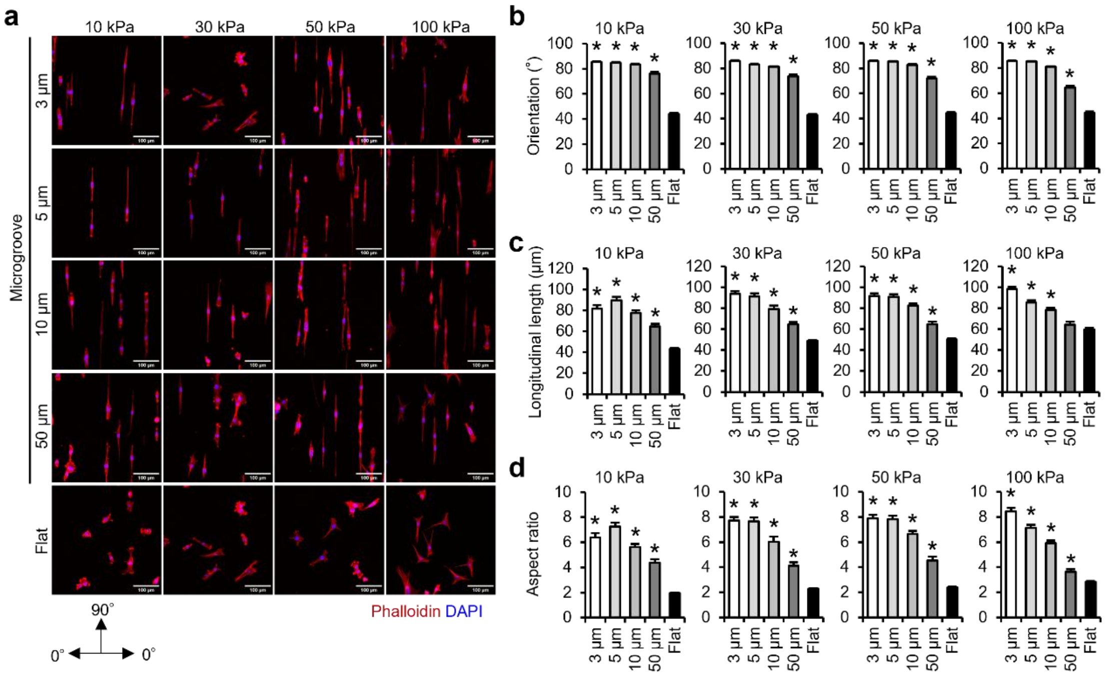


**Sup. Fig. 4 Orientation, aspect ratio, and longitudinal length of myoblasts cultured on a gelatin gel with transferred microgrooves.**

(a) Fluorescence images of myoblasts cultured on 10–100 kPa gelatin gel with 3, 5, 10, and 50 µm microgrooves and flat condition. The images demonstrate myoblasts at 24 h post-seeding, stained for actin (red: phalloidin) and nuclei (blue: DAPI). Scale bars, 100 µm. (b) Orientation of myoblasts. The orientation is analyzed from the longitudinal axis of myoblasts, with the vertical direction of the microgrooves set at 0° and parallel direction at 90°. (c) The longitudinal length of myoblasts is analyzed based on myoblasts stained with phalloidin. (d) Aspect ratio of myoblasts. The aspect ratio is calculated by dividing the length of the longitudinal axis by the length of the transverse axis of myoblasts stained actin. Scale bars, 100 µm. Data are presented as the mean ± S.E.M., n = 25–50 (images), *$p < 0.05$, vs. (Flat) (one-way ANOVA followed by Dunnett's post hoc test).

**Supplementary Table**

**Sup. Tab. 1 Primer sequences for qRT-PCR**

Primer sequences for each gene were used in this study.

| | | |
|---|---|---|
| MYH1 | F | 5'-CCCTAAAGGCAGGCTCTCTCA-3' |
| | R | 5'-GGCGTCAAAAGGCTTGTTCT-3' |
| MYH2 | F | 5'-CTGTGGAATGACCGAAGCGA-3' |
| | R | 5'-TGCAAAGGAACTTGGGCTTTT-3' |
| MYH4 | F | 5'-CATCTGGTAACACAAGAGGTGC-3' |
| | R | 5'-GACTTCCGGAGGTAAGGAGC -3' |
| MYH7 | F | 5'-TACTTGCTACCCTCAGGTGG-3' |
| | R | 5'-ATGGCTGAGCCTTGGATTCTC-3' |
| GLUT4 | F | 5'-GTAACTTCATTGTCGGCATGG-3' |
| | R | 5'-AGCTGAGATCTGGTCAAACG-3' |
| COX IV | F | 5'-GTACCGCATCCAGTTTAACGA-3' |
| | R | 5'-CCATACACATAGCTCTTCTCCCA-3' |
| Myoglobin | F | 5'-TCAAATCTCAGCTGACAGCCA-3' |
| | R | 5'-GGTGAGTCTTAAACAGACCGAT-3' |
| PGC-1α | F | 5'-TGATGTGAATGACTTGGATACAGAC-3' |
| | R | 5'-GCTCATTGTTGTACTGGTTGGATATG-3' |
| GAPDH | F | 5'-GCATCTTCTTGTGCAGTGCC-3' |
| | R | 5'-TACGGCCAAATCCGTTCACA-3' |
| Hprt1 | F | 5'-GTGTTGGATACAGGCCAGACTTTG-3' |
| | R | 5'-GCTGGCCTATAGGCTCATAGTGC-3' |
